# A Wearable Brain–Computer Interface for Mitigating Car Sickness via Attention Shifting

**DOI:** 10.1101/2024.09.25.614936

**Authors:** Jiawei Zhu, Xiaoyu Bao, Qiyun Huang, Tao Wang, Li Huang, Yupeng Han, Haiyun Huang, Junbiao Zhu, Jun Qu, Kendi Li, Di Chen, Ya Jiang, Kailin Xu, Zijian Wang, Wei Wu, Yuanqing Li

**Author notes:** These authors contributed equally to this work (Jiawei Zhu, Xiaoyu Bao). Contributing authors.

## Abstract

Car sickness, an enormous vehicular travel challenge, affects a significant proportion of the population. Pharmacological interventions are limited by adverse side effects, and effective nonpharmacological alternatives remain to be identified. Here, we introduce a novel attention shifting method based on a closed-loop, artificial intelligence (AI)-driven, wearable mindfulness brain–computer interface (BCI) to alleviate car sickness. As the user performs an attentional task, i.e., focusing on breathing as in mindfulness, with a wearable headband, the BCI collects and analyses electroencephalography (EEG) data via a convolutional neural network to assess the user’s mindfulness state and provide real-time audiovisual feedback. This approach might sustainedly shift the user’s attention from physiological discomfort towards the BCI-based mindfulness practices, thereby mitigating car sickness symptoms. The efficacy of the proposed method was rigorously evaluated in two real-world experiments, namely, short and long car rides, with a large cohort of more than 100 participants susceptible to car sickness. Remarkably, over 83% of the participants rated the BCI-based attention shifting as effective, with significant reductions in car sickness severity, particularly in individuals with severe symptoms. Furthermore, EEG data analysis revealed a neurobiological signature of car sickness, which provided mechanistic insights into the efficacy of the BCI-based attention shifting for alleviating car sickness. This study proposed the first large-scale validated, nonpharmacological and wearable intervention method and system for car sickness, with the potential to transform the travel experiences of hundreds of millions of people suffering from car sickness, which also represents a new application of BCI technology.

## 1 Introduction

Car sickness is a typical form of motion sickness that affects passengers in cars. Symptoms can vary in severity, ranging from profuse sweating, mild dizziness and pale complexion to unbearable nausea and vomiting, which prevent passengers from enjoying the convenience of transport [1–3]. Nearly 46% of the global population has experienced car sickness in the last five years. In particular, approximately 12% of adults experience it frequently, and the rate in children is even higher [4, 5]. Therefore, the development of potential treatments for car sickness is highly important.

To date, several valuable neuropsychological theories have been proposed to elucidate the underlying pathological mechanisms of car sickness, thereby guiding the design of treatments. Among these, the widely accepted sensory conflict theory postulates that car sickness arises from a discrepancy between the information received by different sensory modalities and the expectations of the central nervous system [6]. When an individual is exposed to a motion environment, such as being inside a moving vehicle, the vestibular system perceives the actual motion, whereas the visual and/or auditory systems may receive conflicting signals. If the conflict persists or becomes overwhelming, it results in the manifestation of motion sickness symptoms [7]. In recent years, there has also been a growing focus on investigating the neurobiological signatures of motion sickness via electroencephalography (EEG). The evidence converges to indicate an association between the power of the beta oscillation and the severity of motion sickness [8–11]. However, the majority of such studies conducted thus far have focused on motion sickness in general, rather than specifically on car sickness, and have been carried out in simulated indoor environments, lacking rigorous validation in real-world car riding scenarios with a large cohort susceptible to car sickness. Furthermore, previous studies have not yet established an association between the neurobiological signatures of motion sickness, sensory conflict theory, and potential intervention strategies. These factors collectively contribute to a persistent lack of direct evidence establishing a causal link between sensory conflict and car sickness symptoms.

Current treatments for car sickness primarily involve medications such as dimenhydrinate, promethazine, cyclizine, and scopolamine, which have success rates between 62% and 78% [3, 12–14]. However, these moderately effective medications are associated with adverse effects such as headaches, dry mouth, fatigue, dizziness, blurred vision, nausea, and vomiting [12, 13, 15], restricting their use to rare occasions. This has spurred interest in alternative nonpharmaceutical interventions. Multiple intervention approaches have been proposed, such as 1) cue-based methods promoting passengers’ anticipation or comprehension of car motion through additional visual, tactile, or auditory cues [16–21]; 2) attention shifting methods, which switch individuals attention away from discomfort to controlled breathing [22, 23] or cognitive tasks [24]; and 3) biofeedback methods, which are based on physiological signals of heart rate, skin conductance and systolic blood pressure, allowing individuals to self-regulate their physiological state [25, 26]. Among these, attention shifting methods stand out for their simplicity and ease of implementation. However, maintaining sustained attention in dynamic vehicular environments remains challenging due to the continuous sensory conflicts inherent in car rides. Furthermore, the effectiveness of these approaches during real-world car rides remains to be fully established, especially given that most of the previous intervention studies were conducted in simulated laboratory environments, with a small number of participants who were not specifically recruited from populations highly susceptible to car sickness. To our knowledge, there has not been a nonpharmaceutical effective intervention method widely used during everyday car rides.

In this study, we propose a novel attention shifting method based on mindfulness brain–computer interface (BCI) for alleviating car sickness in real-world scenarios. A BCI system establishes a direct interactive control channel between the brain and external devices (such as computers), including modules for brain signal acquisition, brain signal analysis, execution/application, and neurofeedback [27, 28]. By establishing a closed-loop system between neural activity and sensory feedback, BCI can help users achieve and maintain sustained attention [29]. Herein, we employ mindfulness meditation as a specific attentional task, whose core element is to anchor our attention to a specific object, such as breath. It harnesses endogenous top-down control, requiring no external apparatus and remaining robust to vehicular motion [30, 31]. Our approach involves a headband equipped with one channel of electrode automatically receiving and amplifying prefrontal electroencephalography (EEG) signals. The EEG signals are wirelessly transmitted to a computing terminal (e.g., a mobile phone, a tablet PC, or a laptop) and further analysed to assess the users mindfulness state with a convolution neural network (CNN) model. The assessment results are fed back to the user through mindfulness scores and 2D/3D dynamic audiovisual scenes to help users maintain focus on mindfulness meditation (Fig. 1). This closed-loop design enables users to continuously switch attention from physiological discomfort to mindfulness meditation, potentially mitigating sensory conflicts underlying car sickness.

**Fig. 1.**
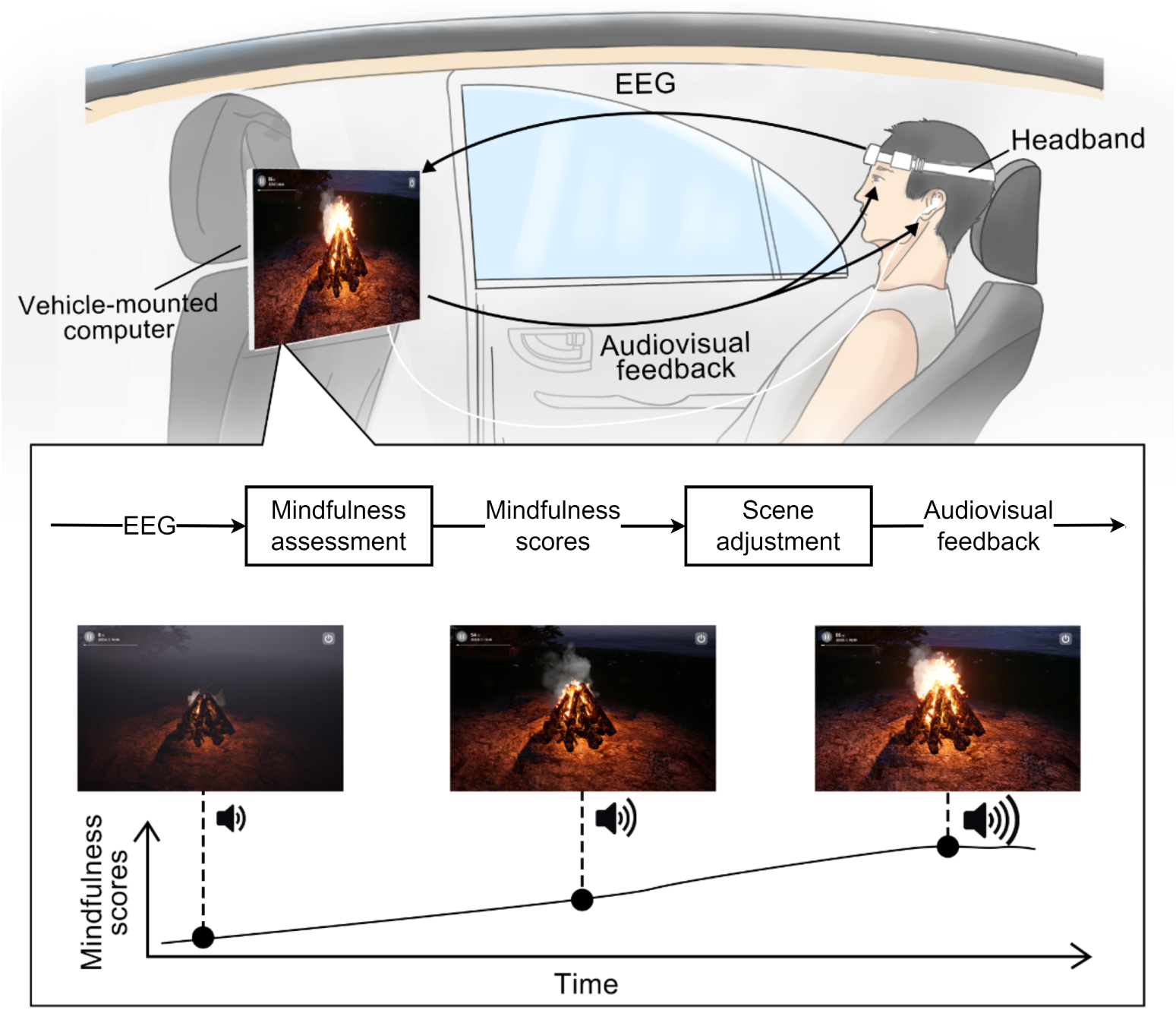
Schematic overview of the proposed mindfulness BCI system. A wireless headband acquires EEG signals, which are transmitted to the computing terminal for real-time assessment of the users mindfulness state on the basis of a pretrained CNN model and generation of mindfulness scores. The mindfulness scores are then used to adjust the context of an interactive audiovisual scene to produce dynamic audiovisual feedback, aiding the user in maintaining sustained attention on mindfulness meditation via self-regulation. For example, the campfire meditation scene displays higher flame intensity and sound volume as mindfulness scores increase.

We conducted two prospective experiments during actual car rides to assess the efficacy of the BCI-based attention shifting for mitigating car sickness. A total of 106 participants prone to car sickness participated in experiment 1 with two sessions, each of which included two consecutive runs of 20-minute car rides on city roads in Guangzhou, China, per session, one with BCI-based mindfulness meditation (mindfulness state) and the other without (control state). The experimental results validated the effectiveness of the BCI-based attention shifting in mitigating car sickness symptoms during short car rides, as evaluated by the Misery Scale (MISC) score [32]. Subsequently, 101 of these participants progressed to experiment 2, which comprised two 120-minute car-riding runs: a mindfulness state and a control state. The results demonstrated the systems effectiveness in the real world with prolonged car rides, which are more prone to inducing severe car sickness symptoms. Furthermore, we analysed the EEG data collected during the two experiments and identified a carsickness-related neurobiological signature, prefrontal beta relative power. Combining the neurobiological signature, attention shift and sensory conflict theory, we provide mechanistic insights into the efficacy of the mindfulness BCI in alleviating car sickness.

## 2 Results

In this section, we present the experimental results regarding mindfulness BCI-based car sickness alleviation and relevant EEG signatures.

### 2.1 BCI-based attention shifting alleviates car sickness in both short and long car rides

Following the completion of experiments 1 and 2, the participants rated the effectiveness of the BCI-based attention shifting for reducing car sickness on a 7-point Likert scale (Fig. 2 (a) and (d)). In experiment 1, 89 out of 106 participants (83.96%) reported that the BCI-based attention shifting effectively alleviated their car sickness during short car rides (Fig. 2(a)). Owing to apprehension about long-duration car rides, five participants voluntarily withdrew from experiment 2. Among the 101 participants, 90 (89.11%) reported that the BCI-based attention shifting was effective in relieving car sickness on long trips (Fig. 2(d)).

**Fig. 2.**
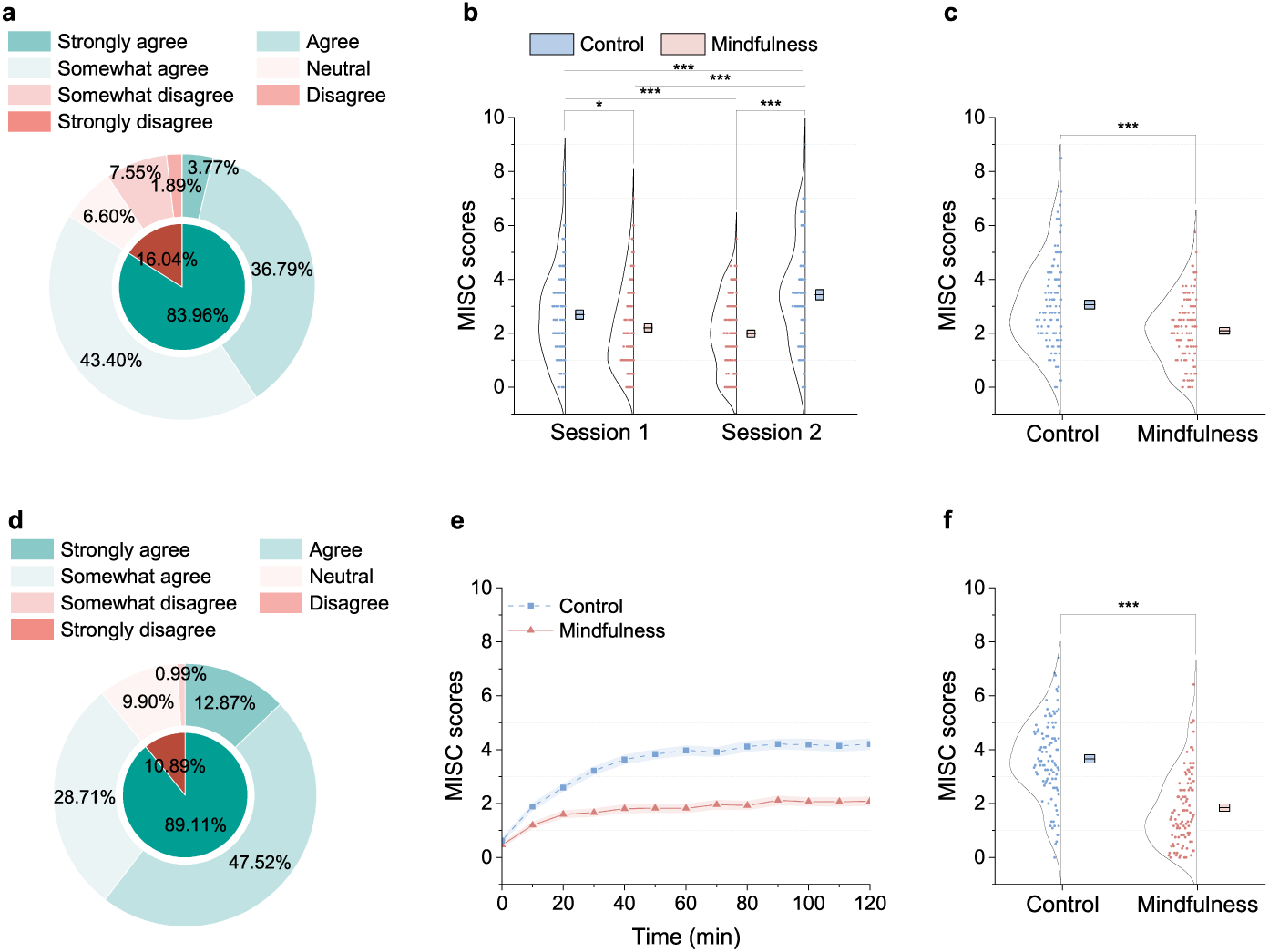
Behavioural results of experiments 1 and 2. Evaluation results for the effectiveness of the BCI-based attention shifting for reducing car sickness during short (a) and long (d) car rides. (b) Two-way ANOVA revealed a significant main effect of state (*F* (1, 420) = 35.731*, P <* 0.001) and a significant main effect of order (*F* (1, 420) = 8.717*, P* = 0.003) but no significant interaction between state and order. Post hoc analyses with Bonferroni correction revealed significantly lower MISC scores in the mindfulness state than in the control state within and across sessions of short-term car rides. (c) The average MISC score across sessions 1 and 2 was significantly greater in the control state than in the mindfulness state (two-tailed, paired *t*(105) = *−*7.230*, P <* 0.001, Cohen’s *d* = *−*0.702). (e) The curves represent the average MISC scores across all participants recorded every 10 minutes in the long car ride, with the red solid line (marked by triangles) representing the mindfulness state and the blue dotted line (marked by squares) representing the control state. The shaded region indicates the standard error of the mean. (f) Scatter plot of the mean MISC scores of each participant across all time points during the long car ride. A significant difference (two-tailed, paired *t*(100) = *−*12.251*, P <* 0.001, Cohen’s *d* = *−*1.219) was observed in the mean MISC scores between the mindfulness state (mean = 1.847, s.e.m. = 0.139) and the control state (mean = 3.659, s.e.m. = 0.151). The boxes represent the standard error of the mean, and the centrelines represent the means. \**P <* 0.05, \*\*\**P <* 0.001.

We then examined the impact of the mindfulness BCI system on the severity of car sickness, as measured with the MISC score. In experiment 1, there was no significant difference in the baseline MISC scores between the two sessions, where the baseline MISC scores were collected at the beginning of each session (two-tailed paired t test, *t*(105) = *−*1.074*, P* = 0.285). Two-way ANOVA revealed a significant main effect of state (mindfulness vs. control: *F* (1, 420) = 35.731*, P <* 0.001) and a significant main effect of order (control state first vs. mindfulness state first: *F* (1, 420) = 8.717*, P* = 0.003), but no interaction between the state and order (*F* (1, 420) = 2.606*, P* = 0.107) (see Fig. 2(b)). Post hoc analyses revealed that the MISC scores in the mindfulness state were significantly lower than those in the control state both within each session (session 1: two-tailed, paired *t*(105) = *−*3.034*, P* = 0.018, Cohen’s *d* = *−*0.295; and session 2: two-tailed, paired *t*(105) = *−*8.574*, P <* 0.001, Cohen’s *d* = *−*0.833; Bonferroni-corrected) and across sessions (session 1 mindfulness state vs. session 2 control state: two-tailed, paired *t*(105) = *−*6.845*, P <* 0.001, Cohen’s *d* = *−*0.665; and session 2 mindfulness state vs. session 1 control state: two-tailed, paired *t*(105) = *−*4.073*, P <* 0.001, Cohen’s *d* = *−*0.396; Bonferroni-corrected). Furthermore, we averaged the MISC scores reported by participants across sessions 1 and 2, and the average MISC score in the mindfulness state was significantly lower than that in the control state (two-tailed, paired *t*(105) = *−*7.230*, P <* 0.001, Cohen’s *d* = *−*0.702; Fig. 2(c)). These results demonstrate the efficacy of the BCI-based attention shifting in alleviating car sickness symptoms during short car rides.

Additionally, the MISC scores in the control state reported during the first 20 minutes of session 1 were significantly lower than those reported during the last 20 minutes of session 2 (two-tailed, paired *t*(105) = *−*5.090*, P <* 0.001, Cohen’s *d* = *−*0.494, Bonferroni-corrected; Fig. 2(b)). This implied that car sickness was less severe in the first 20 minutes than in the last 20 minutes, which is consistent with our daily experience that car sickness symptoms tend to worsen with increased duration of travel. We then examined the impact of the mindfulness BCI system on car sickness symptoms during two 120-minute runs of car rides in experiment 2. Two-way ANOVA was performed to analyse the effects of state (mindfulness state vs. control state) and time (every 10 minutes from zero to the end of each state). A statistically significant interaction was observed between the effects of state and time (*F* (12, 2600) = 7.017*, P <* 0.001). Simple main effects analysis revealed that the MISC scores were significantly affected by both time (*F* (12, 2600) = 39.247*, P <* 0.001) and state (*F* (1, 2600) = 602.289*, P <* 0.001; see Fig. 2(e)). Furthermore, a one-way ANOVA of time within each state revealed that the MISC scores in the mindfulness state (*F* (12, 1300) = 7.869*, P <* 0.001) exhibited a slower growth pattern than those in the control state did (*F* (12, 1300) = 34.999*, P <* 0.001; see Fig. 2(e)). Post hoc analyses revealed that initially, at 0 minutes, there was no difference in the MISC scores between the mindfulness and control states (two-tailed, paired *t*(100) = *−*1.809*, P* = 0.955, Bonferroni-corrected). However, from the 10^th^ minute onwards, the MISC scores in the mindfulness state were significantly lower than those in the control state were (two-tailed, paired t test, *P <* 0.001 every 10 minutes, Bonferroni-corrected). Moreover, when the MISC scores were averaged over time (from 10 minutes to 120 minutes) for each participant, the mindfulness state presented significantly lower average MISC scores than did the control state (two-tailed, paired *t*(100) = *−*12.251*, P <* 0.001, Cohen’s *d* = *−*1.219; Fig. 2(f)). Therefore, from the 10^th^ minute onwards, throughout the two 120-minute car rides, participants reported a significant reduction in the severity of car sickness during the mindfulness state compared with the control state. Moreover, the increasing trend of the MISC score over time was significantly mitigated by the BCI-based attention shifting.

### 2.2 BCI-based attention shifting is more effective for individuals experiencing severe car sickness

We also observed an association between the severity of car sickness in the control state and the effectiveness of the BCI-based attention shifting. After excluding those without clear car sickness symptoms (i.e., MISC score *<* 2 in the control state) during the short (*n* = 26) and long (*n* = 14) car rides, the remaining participants were divided into two groups for both experiments, with a cut-off of 4 for their average MISC scores in the control state (Fig. 3 (a) and (d)). The proportion of participants experiencing relief (lower MISC scores in the mindfulness state) and the magnitude of this reduction increased with car sickness severity in the control state. Specifically, in experiment 1, 72.92% and 100% of the participants in groups 1 and 2, respectively, experienced relief from car sickness symptoms (Fig. 3(b)). Among those participants who experienced relief, the average decreases in MISC scores (control vs. mindfulness) were 1.02 and 2.31 in groups 1 and 2, respectively (Fig. 3(c)). The same pattern was observed as the car ride duration increased to 120 minutes. In experiment 2, 85.71% and 100% of the participants in groups 1 and 2 experienced relief from car sickness symptoms, with average decreases in MISC scores of 1.68 and 2.72, respectively (Fig. 3 (e) and (f)). These results suggest that the BCI-based attention shifting is particularly beneficial for individuals who are more susceptible to car sickness.

**Fig. 3.**
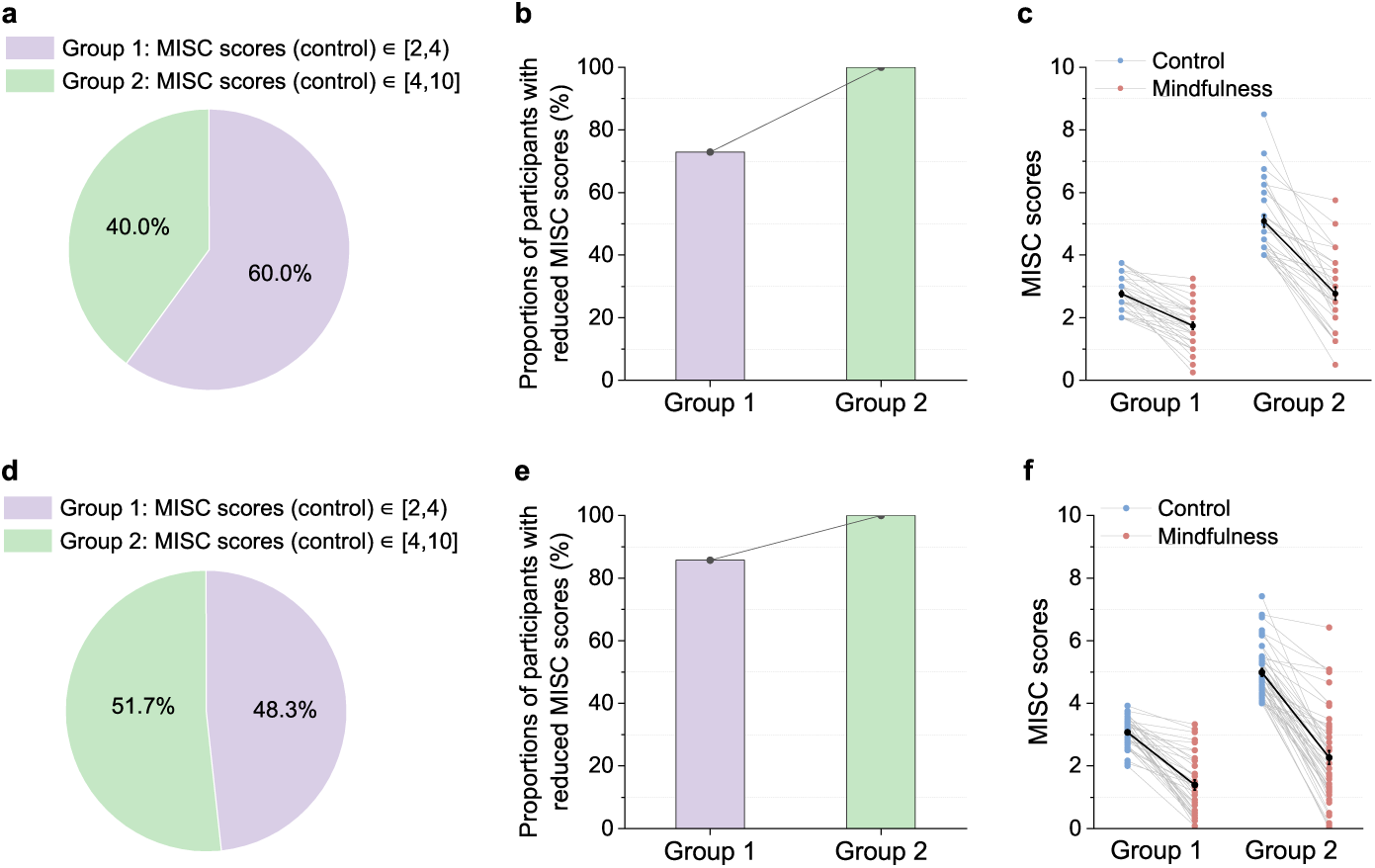
Car sickness relief results for experiments 1 and 2. Proportion of participants in the two groups determined by their average MISC scores in the control state during short (a) and long (d) car rides. Proportions of participants in each group who reported lower MISC scores in the mindfulness state than in the control state during short (b) and long (e) car rides. The MISC scores reported by participants experiencing car sickness relief in the control (blue dots) and mindfulness (red dots) states during short (c) and long (f) car rides. The dark dots represent the group average. All error bars indicate the standard error of the mean.

Furthermore, the BCI-based attention shifting significantly benefited participants experiencing severe nausea (the highest MISC score *≥* 7) in the control state. During the short car ride (experiment 1), these participants (*n* = 18) reported a substantial reduction in symptoms, with average highest MISC scores decreasing from 7.50 to 4.78 and mean MISC scores decreasing from 5.72 to 3.13. Notably, two participants who could not complete both control runs due to extreme discomfort were able to complete the mindfulness runs with only mild dizziness. During the long car rides (experiment 2), a larger group (*n* = 28) with severe nausea experienced similar benefits, with average highest MISC scores decreasing from 7.68 to 4.43 and mean MISC scores decreasing from 5.26 to 2.43. Three participants who had to discontinue the control run early because of severe symptoms successfully completed the mindfulness run without significant discomfort.

### 2.3 EEG signature of car sickness: Prefrontal beta relative power

Beta power, or the ratio of beta power to the entire frequency band, has been preliminarily linked to motion sickness in several previous studies conducted in simulated laboratory environments [8–11]. We analysed the EEG data collected during real-world short car rides (experiment 1) and discovered that the prefrontal beta power ratio relative to the baseline, termed prefrontal beta relative power here, was negatively correlated with changes in the MISC scores relative to the baseline both in the control (Pearson *r* = *−*0.243*, P* = 0.014) and mindfulness (Pearson *r* = *−*0.264*, P* = 0.007) states (Fig. 4 (a) and (b)). Note that the prefrontal beta relative power and MISC score changes were attained by subtracting the corresponding baseline values from the current prefrontal beta power ratio or MISC score, respectively, where the baseline data were collected at the beginning of each run (see Methods for details). Furthermore, the EEG data analysis results of the real-world long car ride (experiment 2) also corroborated this finding in both the control (Pearson *r* = *−*0.207*, P* = 0.042) and mindfulness (Pearson *r* = *−*0.231*, P* = 0.023) states (Fig. 4 (c) and (d)). Notably, four participants in experiment 1 and four participants in experiment 2 were excluded from these analyses because they were extreme outliers (*>* 3 std above or below the mean), resulting in a final sample of *n* = 102 for experiment 1 and *n* = 97 for experiment 2. These results demonstrate that prefrontal beta relative power serves as an EEG signature of car sickness in real-world scenarios.

**Fig. 4.**
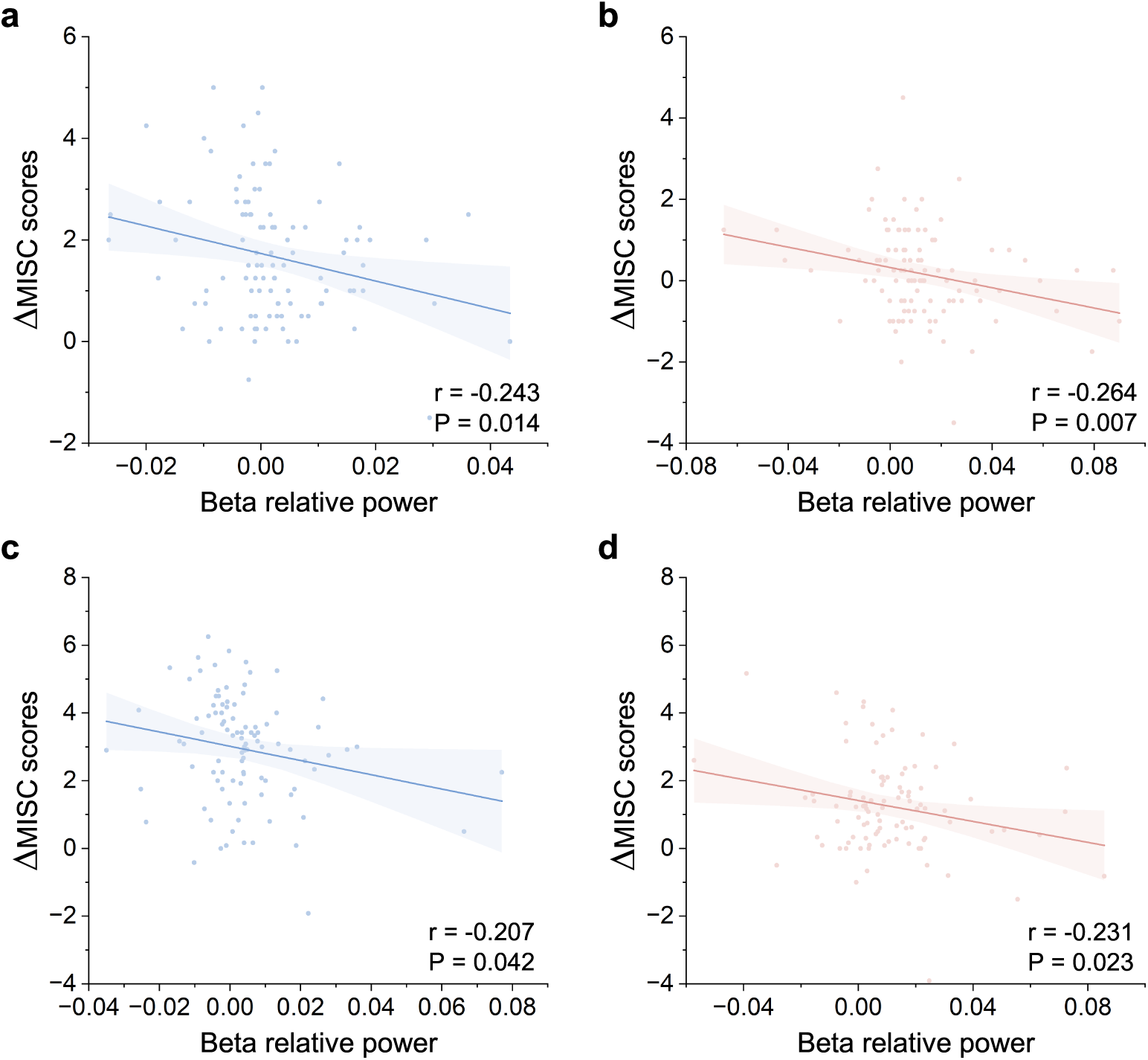
Correlations between prefrontal beta relative power and the relative severity of car sickness in experiments 1 and 2. The MISC scores relative to the baseline scores were negatively correlated with the prefrontal beta relative power in both the control state (Pearson *r* = *−*0.243*, P* = 0.014; (a)) and the BCI-based mindfulness state (Pearson *r* = *−*0.264*, P* = 0.007; (b)) during short car rides. Similar patterns were observed in the control state (Pearson *r* = *−*0.207*, P* = 0.042; (c)) and BCI-based mindfulness state (Pearson *r* = *−*0.231*, P* = 0.023; (d)) during long car rides.

### 2.4 BCI-based attention shifting modulates the EEG signature of car sickness

To further validate the efficacy of the BCI-based attention shifting, we carried out an EEG data analysis to assess its impact on the EEG signature of car sickness, specifically prefrontal beta relative power. As shown in Fig. 5(a), two-way ANOVA revealed a significant main effect of state on prefrontal beta relative power (*F* (1, 404) = 17.851*, P <* 0.001), with no significant main effect of order (*F* (1, 404) = 0.033*, P* = 0.857) or interaction between state and order (*F* (1, 404) = 0.047*, P* = 0.828) in experiment 1. Post hoc analyses indicated that prefrontal beta relative power was significantly greater in the mindfulness state than in the control state, both within (session 1: two-tailed, paired *t*(101) = 3.871*, P <* 0.001, Cohen’s *d* = 0.383; and session 2: two-tailed, paired *t*(101) = 3.220*, P* = 0.007, Cohen’s *d* = 0.319; Bonferroni-corrected) and across sessions (session 1 mindfulness state vs. session 2 control state: two-tailed, paired *t*(101) = 3.073*, P* = 0.011, Cohen’s *d* = 0.304; and session 2 mindfulness state vs. session 1 control state: two-tailed, paired *t*(101) = 3.155*, P* = 0.008, Cohen’s *d* = 0.312; Bonferroni-corrected). Averaging prefrontal beta relative power across sessions for both the control and mindfulness states also confirmed the findings above (two-tailed, paired *t*(101) = 4.502*, P <* 0.001, Cohen’s *d* = 0.446; Fig. 5(b)). Furthermore, the difference in prefrontal beta relative power between the mindfulness and control states was negatively correlated with the difference in MISC relative scores (Pearson *r* = *−*0.324*, P <* 0.001; Fig. 5(c)). In experiment 2, the average pre-frontal beta relative power across time (from 10 minutes to 120 minutes) was also significantly greater in the mindfulness state than in the control state (two-tailed, paired *t*(96) = 3.068*, P* = 0.003, Cohen’s *d* = 0.311; Fig. 5(d)). Moreover, the difference in prefrontal beta relative power between the mindfulness state and the control state was negatively correlated with the difference in MISC relative scores during long car rides (Pearson *r* = *−*0.209*, P* = 0.040; Fig. 5(e)). These results demonstrate that the BCI-based attention shifting modulates the EEG signature of car sickness, providing compelling neurophysiological evidence for its efficacy in reducing car sickness symptoms during both short- and long-term car rides.

**Fig. 5.**
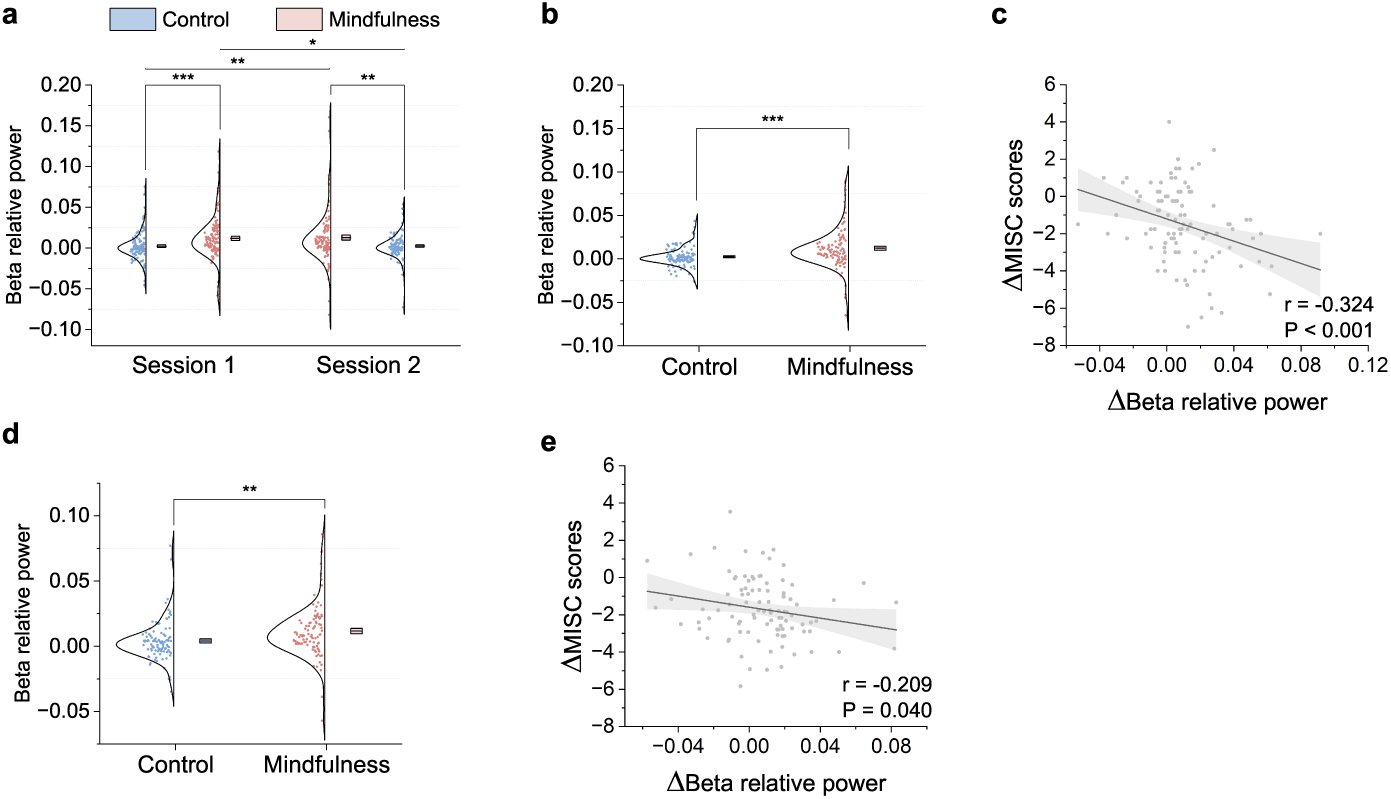
BCI-based attention shifting modulates the prefrontal beta relative power in experiments 1 and 2. (a) Prefrontal beta relative power was significantly lower in the control state than in the mindfulness state in experiment 1 (two-way ANOVA: state: *F* (1, 404) = 17.851*, P <* 0.001). (b) Averaged prefrontal beta relative power across sessions was significantly greater in the mindfulness state than in the control state in experiment 1 (two-tailed, paired *t*(101) = 4.502*, P <* 0.001, Cohen’s *d* = 0.446). (c) The difference in prefrontal beta relative power between the mindfulness state and the control state was negatively correlated with the difference in MISC relative scores in experiment 1 (Pearson *r* = *−*0.324*, P <* 0.001). (d) Averaged prefrontal beta relative power across time was significantly greater in the mindfulness state than in the control state in experiment 2 (two-tailed, paired *t*(96) = 3.068*, P* = 0.003, Cohen’s *d* = 0.311). (e) The difference in prefrontal beta relative power between the mindfulness state and the control state was also negatively correlated with the difference in MISC relative scores in experiment 2 (Pearson *r* = *−*0.209*, P* = 0.040). Boxes and centrelines represent standard errors and means, respectively. \**P <* 0.05, \*\**P <* 0.01, \*\*\**P <* 0.001.

### 2.5 BCI-based attention shifting facilitates attentional engagement

To assess participants’ engagement during BCI-based mindfulness meditation, we examined both the mindfulness scores (derived from the client software) and the theta/alpha power ratio, an established EEG signature of attention [33, 34]. The average mindfulness scores across participants in the mindfulness state runs of experiment 1 and those in the mindfulness state run of experiment 2 are depicted in Fig. 6 (a) and (d), where the mindfulness score curves reach a minimum at the initial moment and every 10-minute timepoint, corresponding to the onset of meditation on the basis of a changed audiovisual scene. The mindfulness scores remained consistently high throughout each 10-minute interval of meditation, indicating that the participants sustained focus on mindfulness meditation with the help of the BCI.

**Fig. 6.**
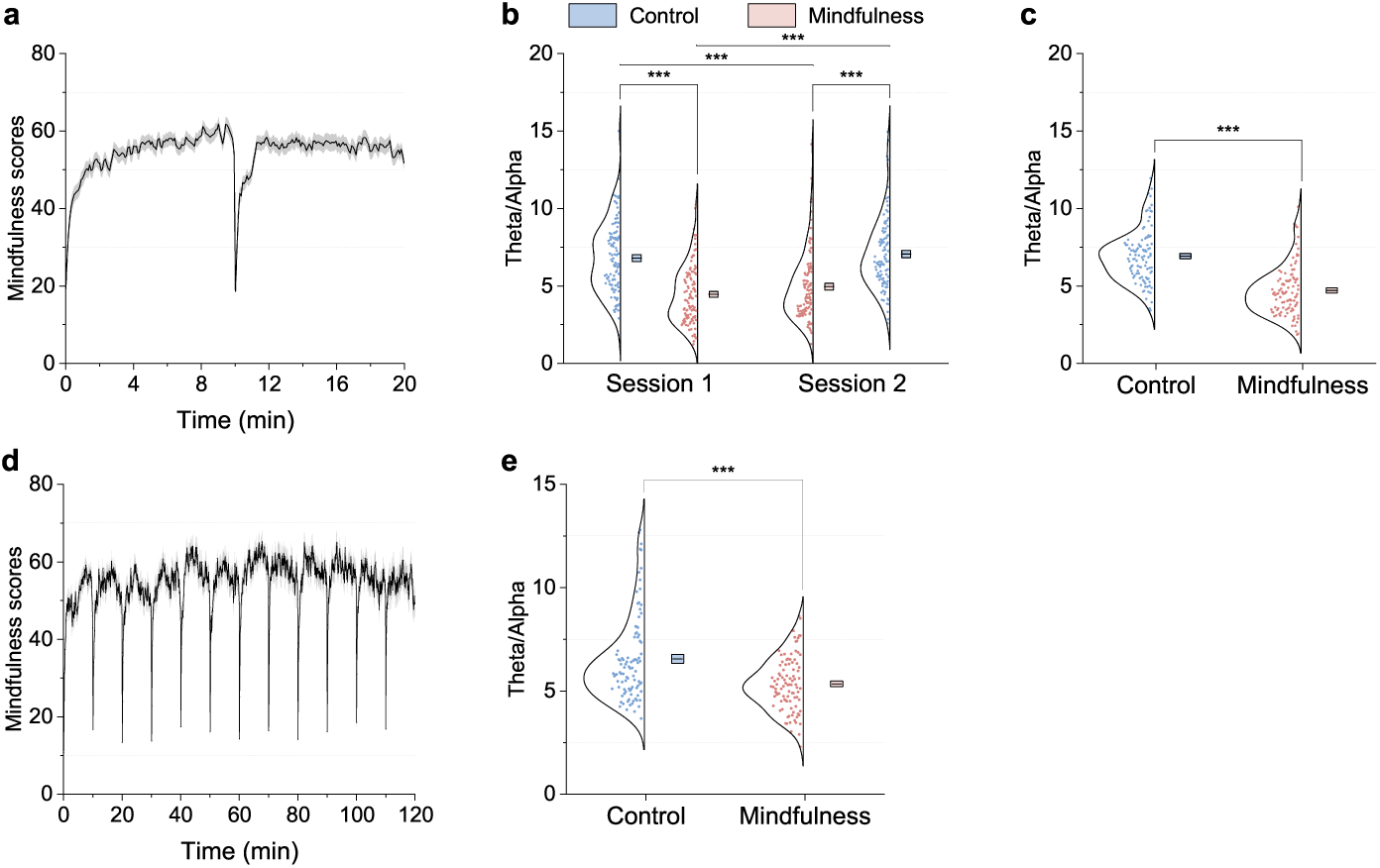
Mindfulness scores and theta/alpha power ratios related to attention in experiments 1 and 2. (a) The mean mindfulness scores of participants in experiment 1 (*n* = 106). The shaded area indicates the standard error of the mean. (b) In experiment 1, the theta/alpha power ratio was significantly lower in the mindfulness state than in the control state (two-way ANOVA: state: *F* (1, 404) = 103.687*, P <* 0.001). (c) The average theta/alpha power ratio in the mindfulness state was significantly lower than that in the control state in experiment 1 (two-tailed, paired *t*(101) = *−*9.822*, P <* 0.001, Cohen’s *d* = *−*0.973). (d) The mean mindfulness scores of participants in experiment 2 (*n* = 101). (e) In experiment 2, the average theta/alpha power ratio during car rides in the mindfulness state was significantly lower than that in the control state (two-tailed, paired *t*(96) = *−*5.064*, P <* 0.001, Cohen’s *d* = *−*0.514). The boxes and centrelines represent standard errors and means, respectively. \*\*\**P <* 0.001.

EEG analysis further supported these findings. In experiment 1, two-way ANOVA revealed a significant main effect of state on the theta/alpha power ratio (*F* (1, 404) = 103.687*, P <* 0.001). Additionally, there was no significant main effect of order (*F* (1, 404) = 0.310*, P* = 0.578) or interaction between state and order (*F* (1, 404) = 3.022*, P* = 0.083) (see Fig. 6(b)). Post hoc analyses confirmed that the theta/alpha power ratio was significantly lower in the mindfulness state than in the control state, both within individual sessions (session 1: two-tailed, paired *t*(101) = *−*8.031*, P <* 0.001, Cohen’s *d* = *−*0.795; and session 2: two-tailed, paired *t*(101) = *−*7.409*, P <* 0.001, Cohen’s *d* = *−*0.734; Bonferroni-corrected) and across sessions (session 1 mindfulness state vs. session 2 control state: two-tailed, paired *t*(101) = *−*8.174*, P <* 0.001, Cohen’s *d* = *−*0.809; and session 2 mindfulness state vs. session 1 control state: two-tailed, paired *t*(101) = *−*5.851*, P <* 0.001, Cohen’s *d* = *−*0.579; Bonferroni-corrected). The theta/alpha power ratios corresponding to the control and mindfulness states were further averaged across two sessions for each participant. As shown in Fig. 6(c), the average theta/alpha power ratio for the mindfulness state was significantly lower than that for the control state in experiment 1 (two-tailed, paired *t*(101) = *−*9.822*, P <* 0.001, Cohen’s *d* = *−*0.973). Similarly, in experiment 2, the average theta/alpha power ratio over time for the mindfulness state was significantly lower than that for the control state (two-tailed, paired *t*(96) = *−*5.064*, P <* 0.001, Cohen’s *d* = *−*0.514; Fig 6e). The reduction in the theta/alpha power ratio, which is indicative of heightened attention [33, 34], provides strong neurophysiological evidence that participants were actively engaged in BCI-based mindfulness meditation during both short- and long-term car rides.

## 3 Discussion

This study introduces a novel attention shifting method based on wearable mindfulness BCI for effectively alleviating car sickness. Eighty-nine of 106 participants (83.96%) in experiment 1 with short car rides (20 minutes) and 90 of the 101 participants (89.11%) in experiment 2 with long car rides (120 minutes) reported that the BCI-based attention shifting effectively alleviated their car sickness. Notably, the BCI-based attention shifting was particularly effective for individuals experiencing severe car sickness, as evidenced by an association between symptom severity and the degree of relief. To rule out potential interference from the placebo effect, a separate experiment was conducted in real-world car riding environments via a sham BCI system with random adjustments to the audiovisual scenes (see the online supplementary material for details). Forty-three participants prone to car sickness were newly recruited to participate in the experiment. The majority of participants (76.74%) reported no car sickness relief from the sham system, with some even experiencing symptom exacerbation (Fig. S1 (a) and (b)), suggesting that the observed benefits of the mindfulness BCI system are attributable primarily to active mindfulness meditation with audiovisual neurofeedback rather than a placebo effect. Furthermore, we demonstrated that the prefrontal beta relative power in EEG exhibited a negative correlation with the relative severity of car sickness and served as a neurobiological signature for car sickness. Importantly, this EEG signature of the prefrontal beta relative power increased when car sickness symptoms were alleviated through BCI-based meditation, which provided a mechanistic explanation for the efficacy of the mindfulness BCI in conjunction with attention shift and sensory conflict theory for car sickness.

Discovering EEG signatures is crucial for understanding the neurobiological mechanism of car sickness. Several prior studies have indicated that beta-band activity in EEG signals might serve as a potential signature for motion sickness, as evidenced by decreased beta-band power in frontal or prefrontal regions associated with heightened motion sickness severity [8–11], which provides valuable cues in identifying neurobiological signatures for car sickness. However, a majority of these studies were carried out in simulated indoor environments and involved a limited number of participants who were not necessarily susceptible to car sickness. Therefore, it remains inconclusive whether beta-band power serves as an EEG signature of car sickness. To address this gap, we conducted experiments for both short and long car rides in a real-world environment with a larger pool of participants prone to car sickness. Our findings reveal that the prefrontal beta relative power negatively correlates with the relative severity of car sickness. This consistent correlation persisted across both the control and BCI-based mindfulness states in both experiments 1 and 2 (Fig. 4). Furthermore, we establish, for the first time, a connection between the EEG signature for car sickness and the intervention method of BCI-based attention shifting. Specifically, the pre-frontal beta relative power recovered when car sickness severity was mitigated through BCI-based mindfulness meditation (Fig. 5). This substantiates our initial discovery that prefrontal beta relative power represents a reliable EEG signature of car sickness.

Sustained attention during mindfulness meditation can be challenging because of its inherent monotonous and intellectually undemanding nature [35, 36], especially within a rapidly changing vehicular environment. To facilitate sustained attention shifting to meditation, the proposed mindfulness BCI system translates EEG-based mindfulness scores into audiovisual neurofeedback, acting as an external monitor of the users’ mindfulness state. This empowers users to endogenously regulate neurofeedback stimuli, helping them effortlessly identify and correct instances of mind wandering during mindfulness meditation, thereby nurturing a deeper and more consistent mindfulness state even during long car rides [37–39]. As shown in Fig. 6 (a) and (d), the participants maintained relatively high mindfulness scores throughout the mindfulness state runs in experiments 1 and 2, indicating the achievement of sustained attention. This observation is further supported by our EEG analysis, which revealed a significant reduction in the theta/alpha power ratio (a well-established neural correlate of attention [33, 34]) during the mindfulness state compared with the control state in experiments 1 and 2 (Fig. 6 (b), (c) and (e)). These results show that participants effectively achieved sustainable attention, shifting from physiological discomfort to BCI-based mindfulness practices.

Combining the EEG signature of car sickness, i.e., prefrontal beta relative power, attention shift and sensory conflict theory, we might offer mechanistic insights into the effectiveness of the mindfulness BCI in alleviating car sickness. According to sensory conflict theory, car sickness might arise when there are disconnects or conflicts between the visual/auditory information perceived and the physical sensations experienced by the body’s proprioceptive and vestibular systems. Using the BCI system, the participants successfully shifted their attention from the changing vehicular environment towards mindfulness practices, thereby reducing sensory conflicts. In this way, their car sickness symptoms were mitigated, which is in line with and extends previous reports on attention shifting methods for alleviating motion sickness, such as controlled breathing [22, 23] and cognitive tasks [24]. Numerous studies have linked beta-band activity to endogenous modulation of cognitive and/or perceptual status [40]. On the one hand, prominent beta-band activity has been observed in tasks involving endogenous, top-down processes [41]. On the other hand, it is well established that in exogenous task settings, which are largely stimulus driven, the appearance of a new sensory stimulus causes a decrease in beta-band activity [42]. In a highly dynamic environment during car rides, the conflicts among various sensory inputs hinder the ability of the brain to effectively synthesize and process the received information, thereby reducing the efficiency of the top-down transfer pathway. Consequently, the brain falls into an exogenously driven mode, which is ultimately reflected in the decrease in the prefrontal beta relative power. We may further postulate that the more sensory conflicts there are, the more severe the symptoms of car sickness, and the lower the prefrontal beta relative power, which was demonstrated in our EEG data analysis results (Fig. 4). Mindfulness meditation, which emphasizes anchoring attention in the present moment or to a specific object such as breath, is a typical type of endogenously driven task [43–45]. When users efficiently shift their attention to sustainable mindfulness meditation via the proposed mindfulness BCI, endogenous regulation of their brain is enhanced, which subsequently leads to an increase in prefrontal beta relative power, as observed in the mindfulness state runs in our experiments (Fig. 5). Taken together, the EEG signature of car sickness and the modulation of this pattern by the BCI-based attention shifting provided further mechanistic insights into the effectiveness of our method for alleviating car sickness.

Previous studies on nonpharmacological interventions for car sickness have been impacted by several methodological challenges. Many studies have been conducted in simulated environments, while few real-world investigations have involved only short car rides (e.g., *<* 20 minutes) and relatively small sample sizes (e.g., *≤* 26 participants) [16, 21, 46, 47]. Prolonged car rides, which are more pragmatic yet arduous to study, have received less attention, although symptoms increase proportionally with exposure duration [1]. Furthermore, in some studies, participants have been recruited without considering their susceptibility to car sickness, and the reported outcomes have often been inconsistent regarding efficacy [17–21]. Consequently, robust, evidence-based nonpharmacological methods for effectively preventing car sickness remain to be established; this deficiency is also reflected in both the current market and the daily experiences of those who suffer from this condition. In this study, we aimed to address these challenges by conducting real-world car-ride experiments with a large cohort of participants specifically selected for their susceptibility to car sickness. These experiments included both short and long car rides, ensuring a comprehensive assessment of the efficacy of the BCI-based attention shifting across varying car ride durations. Our results demonstrate a significant reduction in car sickness symptoms, particularly for individuals experiencing severe symptoms, and are supported by both behavioural and EEG data. This comprehensive evaluation, which is based on a large sample size and real-world conditions, strengthens the validity and generalizability of our findings and holds great promise for alleviating car sickness.

BCIs have promising application prospects in numerous domains, such as motor function assistance and rehabilitation [48], attention training for children with attention deficit hyperactivity disorder (ADHD) [49], depression intervention [50], sleep improvement [51], and awareness detection [52]. For the first time, our work has extended BCI to a new application scenario, namely, alleviating car sickness. Our mindfulness BCI system uses a wearable, cost-efficient headband with dry electrodes for signal acquisition and is compatible with both desktop and mobile terminals, including Windows, Android, and iOS operating systems, without sacrificing effectiveness. This design may facilitate the applicability of the BCI in real-world settings. Furthermore, our mindfulness BCI does not require extensive training, unlike existing BCI systems for intervention or rehabilitation, which may take months to achieve the desired effects. Our mindfulness BCI system with a portable and wearable design enables immediate alleviation of car sickness symptoms and thus provides a readily accessible solution for improving the travel experience of hundreds of millions of people affected by car sickness.

Several important future considerations stemming from the present study are worth noting. Our study focused primarily on the immediate alleviation of car sickness symptoms rather than the long-term therapeutic potential of the BCI-based attention shifting. Future research will be conducted to explore the therapeutic effects of prolonged use of the BCI system on car sickness, i.e., to determine whether symptoms can be permanently alleviated through long-term use of the system. Additionally, while we targeted car sickness in the present study, the potential of the mindfulness BCI system to alleviate other forms of motion sickness, such as seasickness or visually induced motion sickness, warrants further investigation.

In conclusion, we propose a novel attention shifting method based on mindfulness BCI for alleviating car sickness. Rigorous testing in real-world car rides demonstrated that it could effectively mitigate car sickness. With the use of the mindfulness BCI system, most of the participants prone to car sickness experienced either no symptoms or only mild symptoms throughout the entire experiment. Moreover, the experimental results revealed prefrontal beta relative power as a neurobiological marker linked to car sickness. Our findings extend the sensory conflict theory for car sickness and thus offer mechanistic insights into why the BCI-based attention shifting is effective in addressing this condition. To our knowledge, this is the first large-scale validated, wearable, side-effect-free and nonpharmacological tool to alleviate car sickness, which also represents a new practical application of BCI technology.

## 4 Methods

### 4.1 Mindfulness BCI

The mindfulness BCI system consists of three components, including a headband for collecting EEG signals, a computing terminal (e.g., a mobile phone, a tablet PC, or a PC/laptop) with client software for data analysis, and a graphical interface to display audiovisual neurofeedback scenarios. An overview of this system is shown in Fig. 1. We developed a mindfulness BCI system in both desktop and mobile versions with Windows, Android, and iOS operating systems, providing convenience for daily usage. To facilitate data analysis, we used the desktop version in the experiments with a laptop (Windows 11 operating system, Intel i9 13900 CPU, NVIDIA GTX 4060 GPU, and 16 GB RAM). Additionally, the experimental equipment included a pair of headphones and a touchscreen mounted behind the front seat of the car.

A single-channel wearable headband (iHNNK, Inc.) with a sampling rate of 250 Hz was used to acquire EEG signals, whose electrodes could be positioned on either the left or right prefrontal lobe on the basis of the user’s discretion (in this study, the electrode was placed on the right prefrontal lobe), with reference and ground electrodes located at both temples. The electrodes are constructed from hydrogels, and the headband includes a lithium battery, a circuit board for amplifying signals, and Bluetooth for transmitting EEG data wirelessly.

The client software computes mindfulness scores in real time from EEG data, which range from 0 to 100. These scores indicate the mindfulness meditation state of the user, where higher scores indicate a deeper sense of relaxation while maintaining inner focus.

The client software offered different audiovisual scenes developed with Unreal Engine 5.0.3. Each scenario was curated by a professional meditation instructor and included meditation instructions. The audiovisual scenes were adjusted by mindfulness scores, providing real-time audiovisual neurofeedback to the users. Specifically, higher scores enhance the clarity and beauty of scenes and the sharpness of sounds (like bird calls), whereas lower scores result in more indistinct scenes and background sounds, while the volume of the guided instructions remains unchanged. Our experiments utilized four different meditation scenes, i.e., sea wave meditation, campfire meditation, hearing meditation, and desert meditation. The participants could select scenes on the basis of their preference. For example, in the ”campfire meditation” scene, a higher score results in a more intense campfire and louder associated sounds, whereas a lower score diminishes both; participants were instructed to increase the intensity of the campfire’s flames and the volume of its sounds through mindfulness meditation. Users can easily assess their meditation state or notice their mind-wandering through audiovisual feedback, helping them refocus on mindfulness meditation. In general, the participants reported that mindfulness meditation tasks become easier and more enjoyable with the mindfulness BCI system.

### 4.2 Real-time mindfulness meditation assessment

In this study, we propose a cross-subject CNN model for real-time assessment of participants’ mindfulness meditation based on single-channel prefrontal EEG signals.

#### 4.2.1 EEG feature extraction

The single-channel EEG data were segmented into 10-second EEG samples with a 10-second (2500 points) sliding window. For each 10-second EEG sample, feature extraction was carried out as follows. A notch filter with 50 Hz was first applied to the 10-second EEG sample to remove power-line noise. Next, a time–frequency feature matrix was extracted via a filter group of 35 bandpass filters and Hilbert transformation [53]. Specifically, the filter group was applied to the 10-second EEG sample, and 35 filtered 10-second EEG samples with bands from (2*k* + 0.1) Hz to (2*k* + 2.1) Hz were obtained, where *k* = 0*, . . .,* 34. For a specific filtered EEG sample *x*(*t*), Hilbert transformation was first performed:

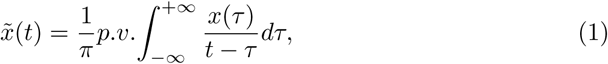

where *x̃*(*t*) was the Hilbert transformation of signal *x*(*t*) and the convolution result of *x*(*t*) and 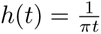, the symbol *p.v.* implies the Cauchy principal value.

Next, the amplitude feature was calculated as follows:

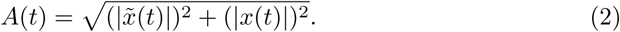

We concatenated the amplitude features of all 35 filtered 10-second EEG samples and obtained a 2-dimensional time–frequency feature matrix with dimensions of 35 *×* 2500 for the 10-second EEG sample. The 2-dimensional feature matrix was downsampled to 35 *×* 100 to reduce the computational cost and avoid overfitting. Furthermore, the feature matrix for the 10-second EEG sample was z score normalized as follows:

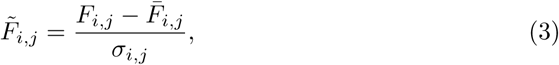

where *F_i,j_*is the element at the *i*^th^ row and the *j*^th^ column of the feature matrix and where *F̄_i,j_* and *σ_i,j_*are the mean and standard deviation, respectively, of *F_i,j_*. A normalized feature matrix with elements *F̃_i,j_* was thus obtained for the 10-second EEG sample. The normalization step was set to avoid undesirable biases and facilitate classification.

#### 4.2.2 CNN model

The cross-subject CNN is shown in Fig. 7, where the extracted time–frequency EEG feature matrix was utilized as its input. The first convolution layer filters the input feature matrix with 32 kernels sized 3 *×* 3 and a stride of 1. Then, a max pooling layer with a pool size of 3 *×* 3 and a stride of 2 follows. The obtained features are fed into the multiscale convolution module, which contains 4 branches with different receptive fields. Notably, to preserve the dimension of the features, convolution and max pooling layers were coupled with padding. Features from different branches were concatenated, sent to a max pooling layer with a pool size of 3 *×* 3 and a stride of 2, and flattened into a vector. The vector was processed by a dropout layer with a drop rate of 0.5 and then fed into a fully connected layer whose kernel size was set to 100. Finally, a Softmax layer was used for classification, and a mindfulness score, the probability of the mindfulness state, was obtained. All the layers except the output layer use the rectified linear unit (ReLU) as the activation function.

**Fig. 7.**
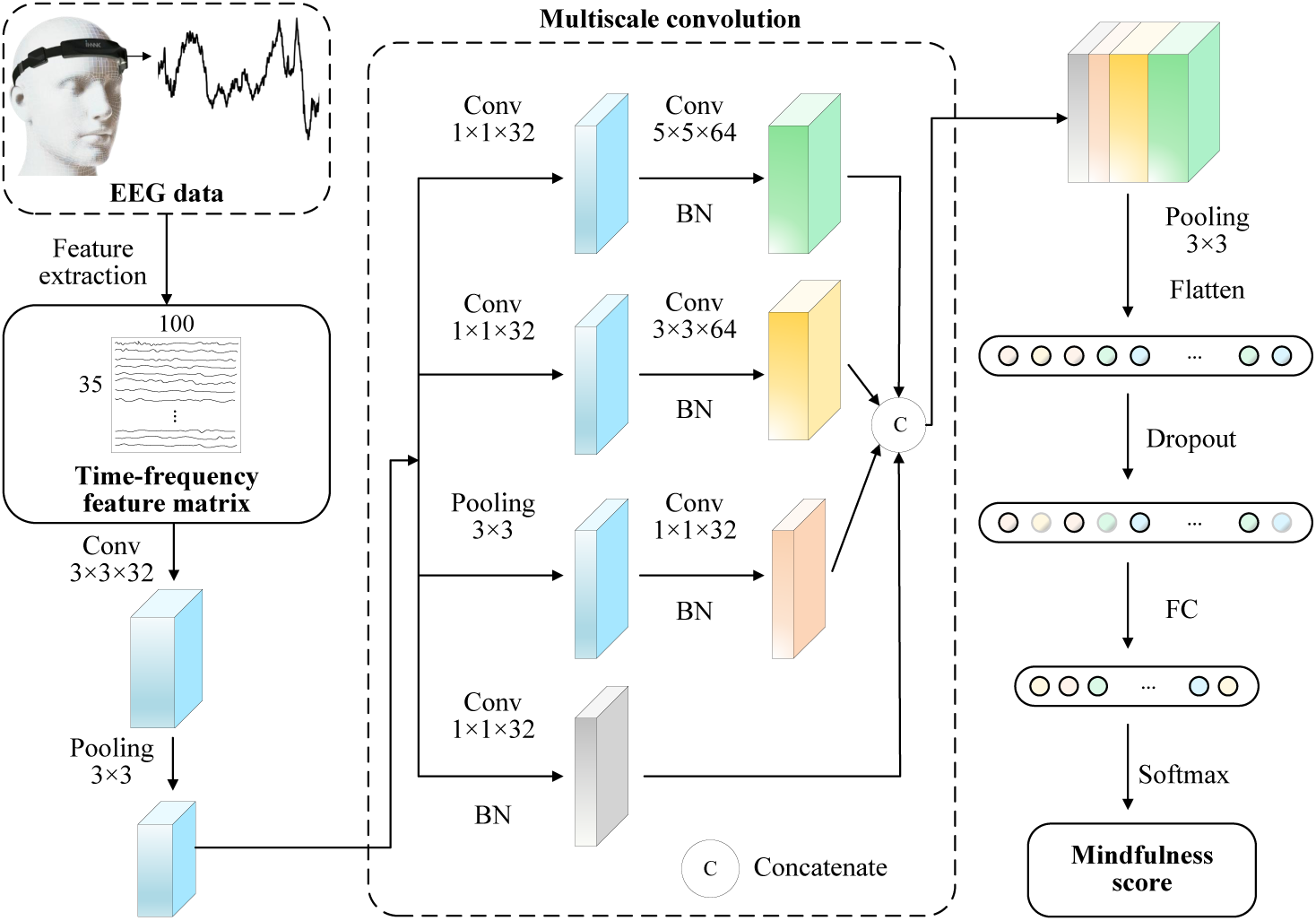
Framework of the cross-subject CNN model used for mindfulness meditation assessment. Conv, Pooling, BN and FC denote the convolution layer, max pooling layer, batch normalization layer and fully connected layer, respectively.

The parameters of the CNN model were initialized with the Xavier initialization method [54], and updated via Adam [55], a method for stochastic optimization, to minimize the cross-entropy loss. The model was trained in a Python environment on an NVIDIA GeForce GPU using a training EEG dataset, which was prepared in our previous studies [56] and included EEG data collected in the mindfulness meditation state and in the rest state. Consequently, the cross-subject CNN model with optimized parameters was obtained.

#### 4.2.3 Online mindfulness meditation assessment

During mindfulness meditation, a subject wore a headband, and the mindfulness BCI system outputs his or her mindfulness scores in real time. Herein, one EEG sample was extracted per second by a 10-second sliding window in real time. For each 10-second EEG sample, a normalized time–frequency feature matrix was calculated as described above and fed into the trained CNN model to output a mindfulness score, which was postprocessed by multiplying it by 100 and presented on the screen. Note that the mindfulness score was updated per second. When the BCI was initially applied to assess mindfulness meditation and output mindfulness scores for a new user, there was not a specific calibration performed at the beginning, which was necessary for most BCI systems; thus, the BCI or the CNN model in this study was cross-subject.

### 4.3 Questionnaires

During the enrolment phase, we used the Motion Sickness Susceptibility Questionnaire shortened version (MSSQ-Short) to estimate the motion sickness susceptibility of the participants [57]. This questionnaire assesses the frequency of occurrence of sickness symptoms across nine different means of transportation and amusement rides, such as cars, coaches, trains, aircraft, funfair rides, etc., and distinguishes between those who have mild motion sickness (percentile scores from 0 to 25%), moderate motion sickness (from 25 to 75%), or severe motion sickness (above 75%). During the experiments, the participants reported their car sickness severity via the MISC (Supplementary Table 1) [32], which measures car sickness symptoms on an 11-point scale; for example, MISC scores of 0, 2, 7 and 10 indicate no problems, vague symptoms, fairly nausea and vomiting, respectively. We built a user-friendly graphic interface for viewing and completing the MISC, displayed on a touch screen attached to the back of the front seat of the vehicle. After experiments 1 and 2, the participants used a 7-point Likert scale, which included strongly agree, agree, somewhat agree, neutral, somewhat disagree, disagree and strongly disagree, to assess the effectiveness of the BCI-based attention shifting for alleviating car sickness.

### 4.4 Experiments

We recruited healthy participants who were susceptible to car sickness through online advertisements and word-of-mouth referrals. A total of 228 individuals signed up for our experiments and reported their susceptibility to motion sickness via the MSSQ-Short [57]. We excluded 122 individuals who never or rarely experienced car sickness during travel, with a susceptibility percentile below 25 according to the MSSQ-Short. The remaining 106 individuals (58 males and 48 females) were included in our study; their mean age was 24.94 years, ranging from 20 to 46 years. These participants frequently experienced car sickness, where their mean MSSQ-Short score was 13.79 (s.e.m. = 0.61), corresponding to a susceptibility percentile of 58.67, indicating moderate susceptibility to car sickness. Furthermore, all the participants reported being in good health; having normal vision and hearing ability; having no history of neurologic, cardiovascular, musculoskeletal, or vestibular impairments; and not being on any medication during the study. Participants were free to withdraw from the study at will and provided informed consent to participate as per procedures approved by the Committee for the Second Affiliated Hospital of South China University of Technology (reference number: 2024-111-01).

Experiments 1 and 2 were designed to explore the car sickness alleviation effect of the mindfulness BCI system under different car riding conditions, as shown in Fig. 1. Experiment 1 consisted of two sessions conducted over two days. In each session, the participants continuously engaged in two runs of short car rides (20 minutes) on city roads in Guangzhou, China: one run with BCI-based mindfulness meditation (mindfulness state) and the other without (control state). The order of the mindfulness and control states alternated between sessions and was counterbalanced among participants. At least one day of rest was provided between the two sessions. During the control state run, participants were instructed to do nothing except to verbally report their car sickness severity via the MISC scale when prompted by an experimenter every two minutes, starting at 0 minutes and continuing until the end of the control state. The experimenter then recorded the rating immediately via a touchscreen. The single-channel prefrontal EEG signals were synchronously recorded. During the mindfulness state run, participants performed BCI-based mindfulness meditation via two 10-minute audiovisual scenes and reported MISC scores at 0, 10, and 20 minutes, which were also recorded by the experimenter. The purpose of reporting MISC scores only three times was to minimize disturbing the participants’ meditation. The participants could selectively focus on changes in sound or audiovisual scenes during BCI-based mindfulness meditation, according to their own preferences. After completing all four runs, the participants were asked to evaluate the effectiveness of the BCI-based attention shifting in alleviating car sickness on a 7-point Likert scale. Over the course of experiment 1, 2 of the 106 participants did not complete the control state car rides due to discomfort.

Experiment 2 consisted of two runs of long car rides, one with BCI-based mindfulness (mindfulness state) and the other without (control state), each lasting 120 minutes on city roads in Guangzhou, China. Five participants voluntarily withdrew from the experiment because of fear of long car rides, leaving 101 participants. The experiment was conducted over two days, with participants completing one run each day. The order of the mindfulness and control states was counterbalanced among the participants. At least one day of rest was provided between the two experimental runs. On one day, participants completed a 120-minute control state run where they did nothing except to report MISC scores every 10 minutes, starting at 0 minutes and continuing until the end of the run. To do this, the participants verbally reported their MISC score, which was then documented by the experimenter using a touchscreen device, while their single-channel prefrontal EEG signals were simultaneously recorded. On the other day, the participants completed a 120-minute mindfulness state run in which they performed BCI-based mindfulness meditation and reported MISC scores every 10 minutes starting from 0 minutes. The experimenter recorded these MISC scores as the laptop ran the mindfulness client software. The participants could selectively focus on changes in sound or audiovisual scenes during BCI-based mindfulness meditation or choose to pause and resume meditating as needed. Notably, if a participant dozed off during either run, the experimenter did not wake them up but allowed them to wake up naturally and then supplement their MISC scores. After completing both runs, the participants evaluated the effectiveness of the BCI-based attention shifting for car sickness alleviation during long car rides on a 7-point Likert scale. Over the course of experiment 2, 3 of the 101 participants did not complete the control state run due to discomfort.

Prior to starting the experiments, we provided the participants the opportunity to familiarize themselves with the mindfulness BCI system, including meditation skills and audiovisual neurofeedback; the duration depended on the mastery of each participant. The recommended meditation skill for users was to focus on their breathing (such as deep breathing and silently counting the number of breaths). Following this advice, the participants were typically able to grasp the usage of the system quickly, usually within 1-3 practice sessions, each lasting 10-20 minutes.

#### 4.4.1 Behavioral data processing

We calculated the percentage of participants who selected positive opinions (i.e., strongly agree, agree and somewhat agree) on the 7-point Likert scale after experiments 1 and 2. The participants’ positive opinions were regarded as approval of the effectiveness of the BCI-based attention shifting for alleviating car sickness.

To compare the differences in the car sickness levels of participants in the control state and the mindfulness state, in experiment 1, the MISC scores reported by each participant at the 10^th^ and 20^th^ minutes of the mindfulness state were averaged as a measure of car sickness severity in the mindfulness state. Similarly, the MISC scores reported at the 10^th^ and 20^th^ minutes of the control state were averaged as a measure of car sickness severity in the control state. In experiment 2, MISC scores reported by participants every 10 minutes in the control state were compared with the MISC scores reported every 10 minutes in the mindfulness state, where each state of each participant consisted of 13 MISC scores, including the MISC score at 0 minutes. For each state, the car sickness relative severity was obtained by subtracting the baseline MISC scores from the average MISC scores, where the baseline MISC scores were reported at 0 minutes in the control or mindfulness state runs. If a participant withdrew from the experiment because of discomfort, their last reported MISC score was used for subsequent analyses.

#### 4.4.2 EEG signature calculation

In experiment 1, to calculate EEG signatures associated with car sickness and attention in the control and mindfulness states, we took three 30-second EEG segments/samples for each participant, one after the MISC scores were reported at 0 minutes of each state and the other two before the MISC scores were reported at the 10^th^ and 20^th^ minutes of each state. Each raw EEG sample was processed, including 1) subtraction of the mean to remove the baseline; 2) notch filtering at 50 Hz; and 3) bandpass filtering using an infinite impulse response filter from 0.5 to 49 Hz. The power spectral density was subsequently estimated from each EEG sample via the Welch method, and several frequency band power values were obtained for each EEG sample, including the power values of theta (4*−*8 Hz), alpha (8*−*13 Hz), beta (13*−*30 Hz), and the entire frequency band (1 *−* 40 Hz). The beta power ratio was denoted as the ratio of the beta power to the entire frequency band for each EEG sample. Next, the prefrontal beta relative power for each of the second and third EEG samples was obtained by subtracting the baseline, the beta power ratio of the EEG sample at 0 minutes, from the corresponding beta power ratio. Finally, the two prefrontal beta relative power values were averaged as the EEG signature of car sickness in each state. Previous studies have shown that the theta/alpha power ratio is related to attention, with a lower frontal theta/alpha power ratio in the meditative state than in the mind-wandering state, and this ratio decreases as the depth of the meditative state increases [33, 34]. Therefore, we calculated two theta/alpha power ratios for the second and third EEG samples and then averaged them as an EEG signature of attention for each state.

In experiment 2, we took a 30-second EEG sample after the MISC scores were reported at 0 minutes and a 30-second EEG sample before the MISC scores were reported every 10 minutes of each state for each participant. The EEG samples corresponding to the period in which the participants doze off were removed. The EEG signatures of car sickness and attention were subsequently calculated as described above. Likewise, for each state, these EEG signatures were averaged across the remaining EEG samples for each participant.

#### 4.4.3 Statistical methods

In experiment 1, we performed a two-way ANOVA to study the impact of two independent variables on each of three dependent variables, where the two independent variables included state (mindfulness state and control state) and order (control state first and then mindfulness state vs. mindfulness state first and then control state), and the three dependent variables included MISC scores, prefrontal beta relative power, and the theta/alpha power ratio. In experiment 2, we used two-way ANOVA to study the impact of state (mindfulness state and control state) and time (every 10 minutes from zero to the end of each state) on the MISC scores. For each dependent variable, when there was a significant difference in the analysis of variance (*P <* 0.05), post hoc comparisons were performed via two-tailed paired t tests with Bonferroni correction for multiple comparisons. For each significant effect, the paired t test effect sizes were measured via Cohen’s *d*. All the statistical analyses were performed via IBM SPSS Statistics 27.0.

## 5 Data availability

The data that support the findings of this study are available from the corresponding authors upon reasonable request.

## 6 Code availability

The code used in the analysis of EEG data reported in this paper is available from the corresponding authors upon reasonable request.

## Acknowledgements

We thank all participants of the study and Zhenfu Wen for valuable help and feedback on the manuscript. This work was supported in part by the STI 2030-Major Projects, China under Grant 2022ZD0208900; in part by the Key Research and Development Program of Guangdong Province, China under Grant 2018B030339001; in part by Guangdong Natural Science Foundation General Program, China under Grant 2024A1515011690; in part by the National Natural Science Foundation of China under Grant 62306120.

## 7 Author contributions

Y.L. was fully responsible for the conceptualization and supervision of this study. Y.L., J.Z. and X.B. designed and organized the experiments. X.B., H.H., J.Z. and J.Q. developed the mindfulness BCI system. J.Z., X.B., Y.H., T.W., L.H., K.L., D.C., Y.J., K.X. and Z.W. collected the data. J.Z., X.B., Q.H. and Y.L. jointly analysed the data. J.Z., X.B., Q.H., W.W. and Y.L. jointly wrote the paper.

## 8 Competing interests

The authors declare that they have no competing interests.

## Notes

### Competing Interest Statement

The authors have declared no competing interest.

### Summary of Updates

Discussion and Methods have been updated.

