## Supplementary Materials for "A Wearable Brain–Computer Interface for Mitigating Car Sickness via Attention Shifting"

### Methods

To assess the potential contribution of the placebo effect to the alleviation of car sickness symptoms, we conducted an additional short-term car riding experiment utilizing a placebo BCI system.

#### Placebo BCI system with sham audiovisual feedback

The placebo BCI system was similar to the proposed mindfulness BCI system, consisting of three components: a headband for EEG data collection, a computing terminal with client software, and a graphical interface displaying audiovisual feedback scenes. These scenes visually resembled those in the mindfulness BCI system but lacked meditation instructions. Importantly, the mindfulness scores were randomly generated and used to adjust the audiovisual feedback scenes every second. Therefore, the placebo BCI system provided sham audiovisual feedback to participants.

#### Placebo experiment

An independent cohort of 43 healthy participants (20 males and 23 females; mean age = 23.67 years, range 20-27) were recruited. The participants reported frequent car sickness, with a mean MSSQ-Short score of 14.72 (s.e.m = 2.59), corresponding to a susceptibility percentile of 61.69, indicating moderate susceptibility to car sickness. There was no significant difference in MSSQ-Short scores between the participants in this placebo experiment and those in experiment 1 (mindfulness BCI vs. placebo; two-tailed $t(147) = 0.706$, $P = 0.481$). Participants were free to withdraw from the placebo experiment at will and provided informed consent to participate.

Prior to the placebo experiment, participants received the same amount of training and instructions on the mindfulness BCI system as the other cohort in this study. However, they were not informed that the feedback they received during the placebo experiment was a sham feedback. The experimental paradigm was identical to experiment 1, with two sessions conducted over two days. Each session involved two consecutive short-term (20 minutes) runs of car rides on city roads in Guangzhou, China, one run with the placebo BCI intervention (placebo state), and the other without (control state). The order of the placebo and control states was alternated between sessions and counterbalanced across participants. A minimum one day rest period was mandated between sessions. During the placebo state run, participants watched two 10-minute audiovisual scenes with sham feedback and reported MISC scores at 0, $10^{th}$ and $20^{th}$ minutes, recorded by an experimenter. Participants could choose to focus on changes in sound or audiovisual scenes based on their own preference. During the control state run, participants were instructed to do nothing except to verbally report their car sickness severity via the MISC scale when prompted by an experimenter every two minutes, starting at 0 minutes and continuing until the end of the control state. The experimenter then recorded the rating immediately. Upon completing the placebo experiment, participants rated the effectiveness of the placebo BCI system with sham audiovisual feedback in reducing car sickness based on a 7-point Likert scale.

**Behavioral data analysis**

The behavioral data processing followed the identical procedure as in experiment 1. Participants' negative opinions (i.e., strongly disagree, disagree, somewhat disagree and neutral) were considered a rejection of the effectiveness of the placebo BCI system in car sickness alleviation. Additionally, for each participant, the MISC scores reported at the 10$^{th}$ and 20$^{th}$ minutes of each session were averaged to measure the severity of car sickness in the control or placebo state. These MISC scores were further separately averaged across sessions 1 and 2 to represent the overall car sickness severity of the control and placebo states, respectively.

Furthermore, we conducted a two-way ANOVA to investigate the impact of state (control state and placebo state) and order (session 1: control state first vs. session 2: placebo state first) on MISC scores. When a significant effect was found ($P<0.05$), post hoc comparisons were performed using two-tailed paired T-tests, with Bonferroni multiple comparison correction. For each significant effect, the effect size of paired T tests was measured using Cohen's d.

### Results

As shown in Fig. S1(a), out of the 43 participants, 33 (76.74%) reported that the placebo BCI with sham audiovisual feedback provided no relief for car sickness and even exacerbated symptoms.

We next examined the impact of the sham audiovisual feedback on car sickness severity. A two-way ANOVA revealed a significant interaction between state (placebo vs. control) and order (control state first vs. placebo state first; $F(1,168) = 8.478$, $P = 0.004$) as well as a significant main effect of order ($F(1,168) = 6.183$, $P = 0.014$). However, no significant main effect of state was observed ($F(1,168) = 0.759$, $P = 0.385$) (see Fig. S1(b)). Post hoc analyses showed that the MISC scores in the placebo state during the last 20 minutes in session 1 were significantly higher than those in the control state during the first 20 minutes in session 1 (two-tailed, paired $t(42) = 3.471$, $P = 0.007$, Cohen's $d = 0.529$; Bonferroni-corrected), and significantly higher than those in the placebo state during the first 20 minutes of session 2 (two-tailed, paired $t(42) = 3.837$, $P = 0.002$, Cohen's $d = 0.585$; Bonferroni-corrected). This suggests that the placebo BCI system with sham audiovisual feedback tended to exacerbate car sickness symptom with prolonged car rides. Additionally, there was no significant difference in the MISC scores between control and placebo states within session 2 (two-tailed, paired $t(42) = 1.823$, $P = 0.453$; Bonferroni-corrected) or across sessions (session 1 placebo state vs. session 2 control state, two-tailed, paired $t(42) = 2.639$, $P = 0.070$; and session 2 placebo state vs. session 1 control state, two-tailed, paired t(42) = 1.664, P = 0.621; Bonferroni-corrected). Furthermore, the averaged MISC scores between the control and placebo states showed no significant difference (two-tailed, paired t(42) = 1.223, P = 0.228, Fig. S1(c)). These findings show negligible or even negative effects of the sham feedback on car sickness, indicating that the observed reduction in car sickness symptoms from the proposed mindfulness BCI was not due to the placebo effect.

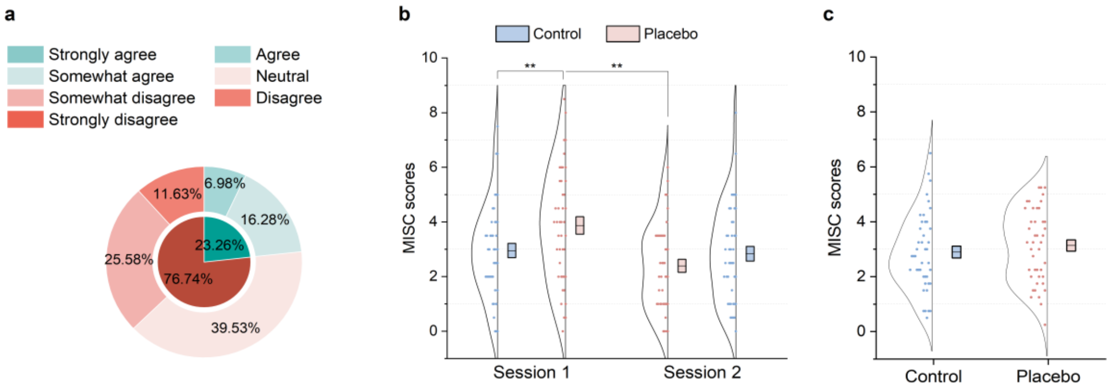


**Figure S1**. Behavioral results from the placebo experiment. (a) Evaluation of the effectiveness of the placebo BCI system with sham audiovisual feedback in alleviating car sickness. (b) Two-way ANOVA revealed a significant interaction between state and order ($F(1,168) = 8.478$, $P = 0.004$), and a significant main effect of order ($F(1,168) = 6.183$, $P = 0.014$), but no main effect of state. (c) Averaged MISC score across sessions 1 and 2, showing no significant difference between the control state and the placebo state (two-tailed, paired $t(42) = 1.223$, $P = 0.228$). Boxes represent the standard error of the mean, and center lines represent the means. $^{**}P < 0.01$.

**Table S1** The MISC used to measure the car sickness severity.

| Symptoms | | MISC |
|---|---|---|
| No problems | | 0 |
| Uneasiness (no typical symptoms) | | 1 |
| Dizziness, warmth, headache, stomach, awareness, sweating, and other symptoms, but no nausea | Vague | 2 |
| | Slight | 3 |
| | Fairly | 4 |
| | Severe | 5 |
| Nausea | Slight | 6 |
| | Fairly | 7 |
| | Severe | 8 |
| | Retching | 9 |
| Vomiting | | 10 |
